# The relationship between varicella (chickenpox) and scarlet fever in contemporary Hong Kong

**DOI:** 10.1101/221507

**Authors:** S. Coleman

## Abstract

Scarlet fever epidemics have reemerged in China, the UK, and Hong Kong. This research tests whether scarlet fever epidemics in Hong Kong are linked to varicella epidemics. Varicella infection is a known risk for invasive Group A Streptococcal infections, and historical research shows a connection between varicella and scarlet fever. This analysis examines the relationship between these two disease in Hong Kong from 2011 to 2015 and compares varicella rates before and after the reintroduction of scarlet fever. Analysis shows that scarlet fever and varicella have synchronous annual epidemic cycles, and a mathematical model of the relationship between scarlet fever and varicella is estimated. Varicella rates were unchanged by the return of scarlet fever, but annual varicella cycles may have influenced the size and timing of scarlet fever outbreaks. Vaccination policies for varicella may need to be adjusted to limit scarlet fever epidemics.

## Introduction

This research examines whether recent scarlet fever epidemics in Hong Kong may be linked to varicella. Varicella is a known risk factor for invasive Group A Streptococcal (GAS) infections, such as toxic shock syndrome and necrotizing fasciitis.^1^ The possibility of an association between scarlet fever (a GAS infection) and varicella is also indicated by a review of historical case studies linking scarlet fever to varicella in the late 19^th^ century and by statistical analysis that shows a strong association between the two diseases in the early 20^th^ century in four American cities—Boston, Chicago, New York, and Philadelphia. The likelihood of a connection between the two diseases is increased because the two diseases attack children in about the same age range, from four to ten years old. As scarlet fever epidemics have unexpectedly returned in several countries, including China^3,4^ and the UK^5^, the question is whether there is once again a connection between varicella and scarlet fever. Prior research does not give a definitive answer, however, as to the nature of this relationship; for example, there may be a causal link or a propensity to co-infections or both. The possibility of a relationship also depends on the vaccination rate for varicella.

## Methods

In this analysis the association between varicella and scarlet fever is investigated in the period 2011 to 2015 in Hong Kong using monthly reported cases of the two diseases. Monthly data is available in English from The Centre for Health Protection, Department of Health, Hong Kong.^6^ All data is aggregate data in the public domain. During this period and earlier, Hong Kong had annual cycles of varicella outbreaks. And, beginning in 2011, Hong Kong also had annual scarlet fever outbreaks. Although it would be desirable to test for a scarlet fever-varicella connection in China and the UK, this is not possible because varicella is not a reportable disease in those countries. The analysis is limited to what can be learned from aggregate data, as data on individual cases is not available

This research concerns two questions: the relationship between the two diseases, and the causal direction, if that can be determined. The analysis begins with a comparison of their monthly time series of reported cases from 2011 to 2015. Following that, the relationship of scarlet fever to varicella is estimated with ordinary least-squares regression. The question of causality is further investigated with a pre/post comparison of the varicella rates before the scarlet fever epidemics began (2009 and 2010) and after (2011 to 2015).

The regression model for the relationship between scarlet fever cases (SF) and varicella cases (V) and time (T) is

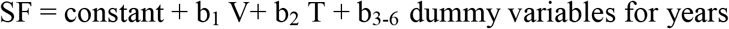

The time variable T is measured in months from the beginning of the time series and allows an estimation of a possible trend in scarlet fever cases. The dummy variables allow inclusion of annual changes in the magnitude of scarlet fever epidemics (for unknown causes). Because of autocorrelation and heteroskedasticity in the scarlet fever time series, a special econometric program, Gretl^7^, was used to estimate a robust standard error for each coefficient to correct for these problems.

The main research question in the pre/post analysis is whether the emergence of scarlet fever in 2011 changed the ongoing pattern of annual varicella cycles. Or did the emerging scarlet fever epidemics develop in correspondence to the pre-existing annual varicella cycles?

## Results

The first part of the analysis concerns the years from 2011 to 2015 when scarlet fever reemerged on a larger scale. Preliminary regression analysis showed that scarlet fever cases in January and February of 2011 were outliers in the model. That is, these two months belonged in the pre-scarlet fever era. So the analysis of scarlet fever begins with March 2011. In this period the average number of scarlet fever cases per month was 112 and for varicella 809. The correlation between varicella cases and scarlet fever cases was 0.59 (p<.0001). The number of annual varicella cases ranged from 650 in 2014 to 1136 in 2011, while scarlet fever cases ranged from 92 in 2013 to 127 in 2011. By contrast, in 2010 there were only 11 reported scarlet fever cases. A comparison of monthly time series plots shows that both diseases have bimodal distributions with peaks, on average, in June and December. Figure 1 shows the average monthly cases for each disease from March, 2011 to December, 2015 expressed logarithmically. (Because there are many more varicella cases than scarlet fever cases, logarithms allow better scaling of the graph for visual interpretation.) This is the same bimodal seasonal pattern of scarlet fever reported in China^4^. A cross-correlation analysis of the monthly time series of both diseases from 2011 to 2015 finds the highest correlation between the two time series at zero lag. That is, the two time series are approximately synchronous at the time-scale of months, but it is not possible to determine if one disease tends to precede the other at a shorter time interval. The incubation and infectious periods of both diseases are shorter than a month.

**Figure 1.**
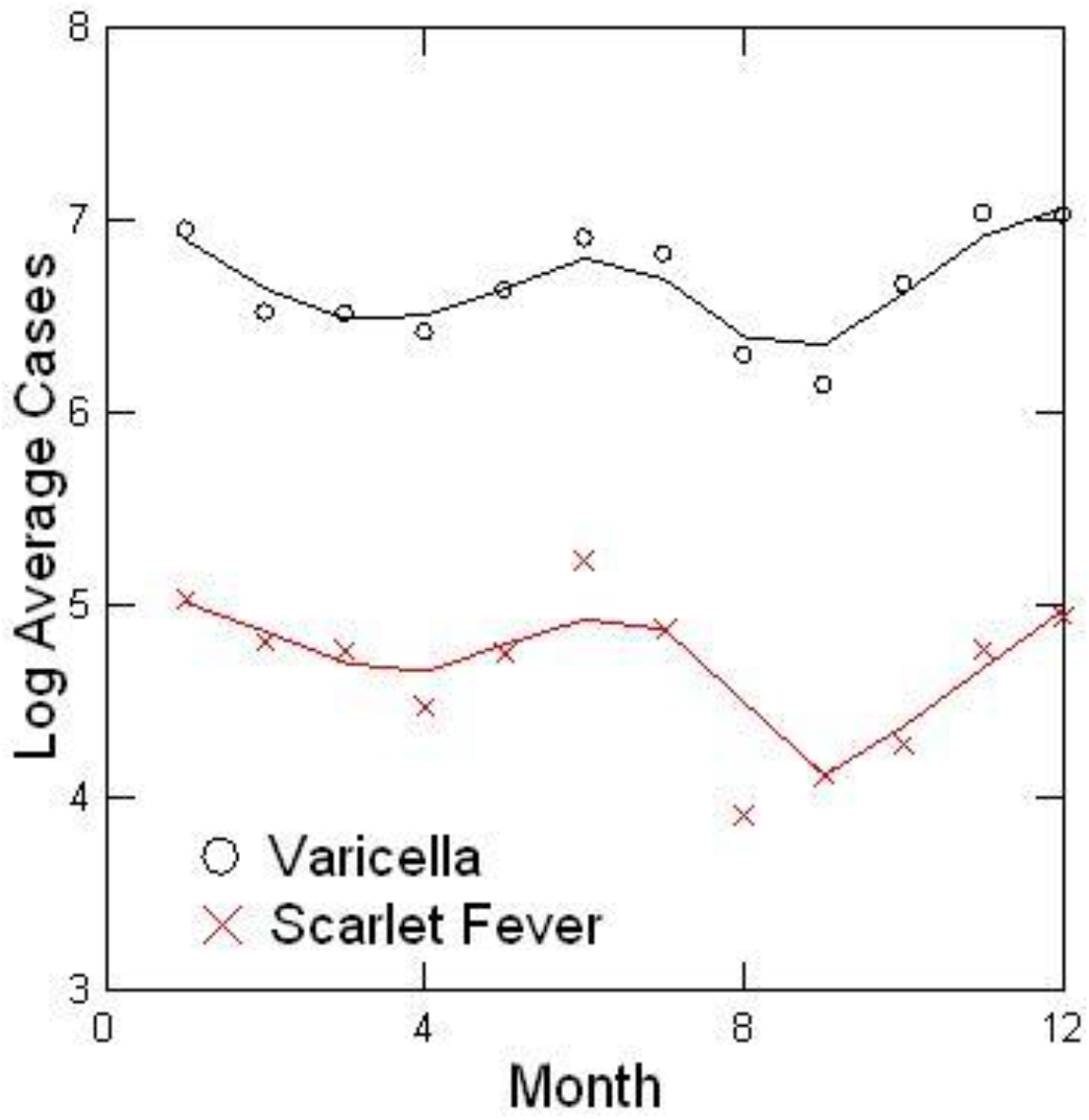
Graph of log average cases per month for scarlet fever and varicella with LOWESS smoothing from March, 2011 to December, 2015.

Estimation of the proposed model shows a moderately strong linear association between varicella and scarlet fever. A comparison of the observed and predicted (estimated) scarlet fever cases is in Figure 2. R^2^ for the estimated model is 0.54 with N = 58. The estimated model (with robust standard errors in parentheses) is

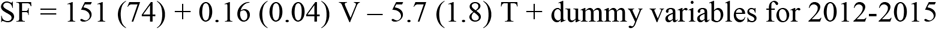

**Figure 2.**
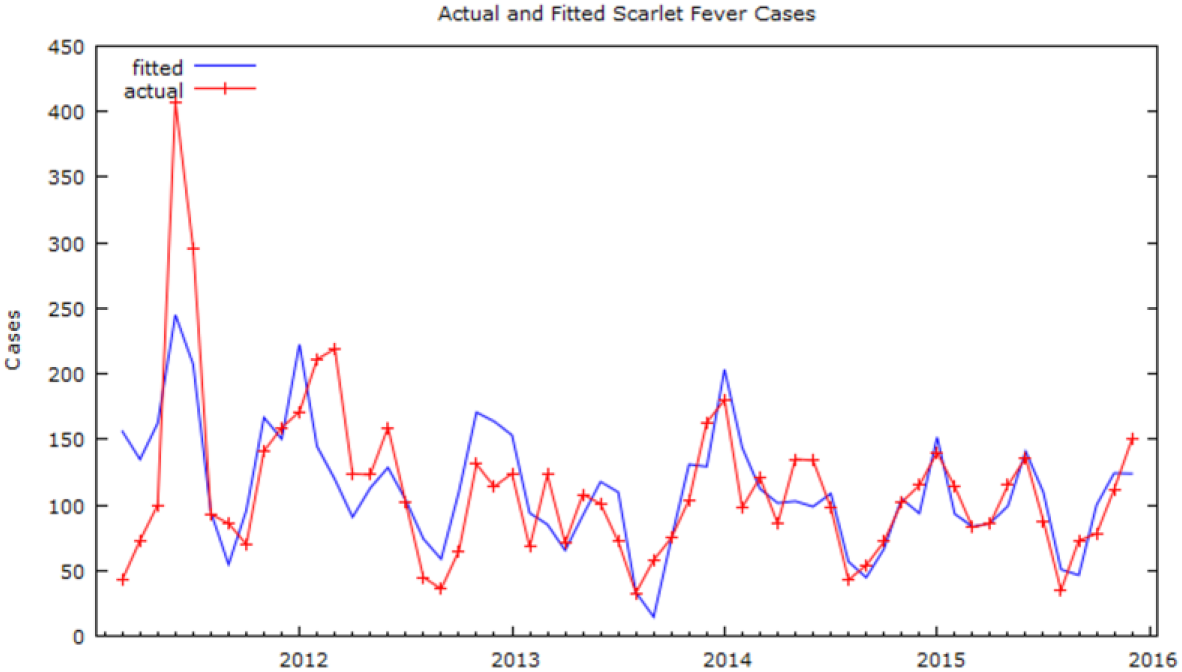
Observed scarlet fever cases by month and fitted (estimated) regression model: March, 2011 to December, 2015.

For every 100 cases of varicella, there are about 16 cases of scarlet fever with 95% confidence interval [8, 24]. The significance level for V is p <.0001; for T, p =.003, indicating a decreasing trend in scarlet fever, after controlling for annual changes. All dummy variables for years are positive and statistically significant (p <.01); that is, there are substantial year-to-year changes in average scarlet fever counts independently of varicella and the decreasing trend. A varicella term with one month lag was tried in the model, but it was not statistically significant.

In the pre/post scarlet fever analysis, the time period is divided at February 2011, which ends the pre-scarlet fever period. Statistical comparison of varicella case numbers in the two periods is based on a t-test of means and a Kolmogorov-Smirnov (K-S) nonparametric test of inequality between cumulative distributions. The tests show no statistically significant difference between mean varicella cases per month in the pre- and post-scarlet fever periods (t-test t = 0.04, difference in means 2.8, p =.97), and no difference in the pre/post cumulative distributions (K-S test, p = 0.76). There also does not seem to be any change in the bimodal seasonality of varicella, which goes back additional years. This means that scarlet fever epidemics did not have a significant impact on varicella rates.

## Discussion

The Hong Kong data captures one of the rare times in history when scarlet fever epidemics arose during a period of ongoing annual varicella cycles. From this analysis one can reject the hypothesis that the association of the two diseases resulted from the influence of scarlet fever on varicella. It seems more likely that varicella influenced the timing and number of scarlet fever cases. But to go further in explaining the connection between the diseases would require data at the individual level.

One might ask whether the lack of routine childhood vaccination for varicella contributed in part to the return of scarlet fever. In the years of this analysis Hong Kong did not routinely vaccinate children for varicella, but in 2013 Hong Kong adopted a plan for vaccination in two doses with the first at age one. The plan did not, however, include catching up older children on vaccinations^8^. The new vaccination program is unlikely to have affected this analysis, however, because varicella usually affects children over the age of three. In China, vaccination for varicella is optional and often only uses one dose, limiting its effectiveness.^9^ A practical question is whether a higher vaccination rate for varicella would inhibit scarlet fever epidemics in Hong Kong and other countries, such as the UK and China, that do not require varicella vaccination.

